# Pleasant smells: a privileged gateway to soothing autonomic responses and improving brain-body rhythm coupling

**DOI:** 10.1101/2024.11.29.625994

**Authors:** Valentin Ghibaudo, Matthias Turrel, Jules Granget, Samuel Garcia, Jane Plailly, Nathalie Buonviso

## Abstract

Aromatherapy commonly uses odors to improve well-being through their evocation of positive emotions. Although knowledge in this area is often very empirical, the olfactory stimulus has different properties which, taken together, could explain why it can relax. First, olfactory sense have a direct access to the limbic system, without thalamic relay processing, which confers it a strong emotional valence. Second, when appreciated, odors can slow down breathing and cardiac rates. Third, when slow and deep, breathing can entrain brain activity, due to the mechano-sensitivity of olfactory receptors to airflows. We hypothesized that, thanks to these properties, pleasant odors could enhance the subjective feeling of relaxation, slow down body rhythms, and facilitate entrainment of brain activity by respiration. Comparing the effects of a personally pleasant odor to a personally pleasant music on psychological, physiological and neuronal responses, we showed a tendency for both odors and music to enhance subjective relaxation. However, only pleasant odors were able to 1) decrease heart rate while increasing its variability, and 2) decrease respiratory rate while enhancing the respiratory drive of brain activities, regardless of the music tempo. Overall, we demonstrated that the positive emotion evoked by a personally pleasant smell is sufficient to evoke an olfactomotor response, which, by slowing breathing, synchronizes respiration, fluctuations of heart rate and brain activity.

## 1 INTRODUCTION

At a time when non-pharmacological interventions are increasingly advocated, the treatment of emotional disorders lends itself particularly well to this type of approach. To achieve positive emotions such as relaxation, humans have long used breath control as a means of influencing the functioning of the autonomous nervous system and the subjective feeling of well-being (Zaccaro et al., 2018). Most of empirically known methods for improving relaxation involve slowing down breathing, which reduces stress effects (Brown & Gerbarg, 2009; Jerath et al., 2015; Zaccaro et al., 2018). Physiologically speaking, slow breathing induces a shift from sympathetic dominance toward a net increase in parasympathetic tone, resulting in heart rate slowdown (Brown & Gerbarg, 2009). At the central level, studies on rodents recently showed that a slow respiratory rhythm, which is associated with a rest state, is also a potent mechanism for modulating brainwave activity (Girin et al., 2021; Hammer et al., 2021; Juventin et al., 2023) and allows large brain networks to synchronize with breathing (Girin et al., 2021; Juventin et al., 2023). This is primarily because slow breathing, when nasal, is optimal to stimulate olfactory receptor cells in the nose, known to be sensitive to air pressure (Grosmaitre et al., 2007). Cognitive factors, such as attention toward breathing or volitional control, also increase coherence between brain activity and breath rhythm in the anterior cingulate, premotor, insular, and hippocampal cortices (Herrero et al., 2018). It is interesting that respiration can modulate brain activity in areas involved in emotion (and various cognitive functions), in both rodents and humans (Folschweiller & Sauer, 2021; Herrero et al., 2018; Zelano et al., 2016).

Conversely, emotions modulate breathing. While anxiety and stress speed up breathing frequency (Boiten et al., 1994; Hegoburu et al., 2011; Masaoka & Homma, 2001), pleasant feelings slow it down (Masaoka et al., 2005). Such an influence of emotions on breathing is based on the important reciprocal connections between amygdala nuclei and respiratory regions (Fulwiler & Saper, 1984). As proof, electrical stimulation of the amygdala accelerates respiratory rate in animals (Harper et al., 1984) or induces apnea in humans (Dlouhy et al., 2015; Lacuey et al., 2019; Rhone et al., 2020).

Olfaction lies at the heart of the close link between emotion and respiration, which leads us to present this whole as a triad of Respiration – Olfaction – Emotions. Odors are an interesting tool to achieve relaxation by slowing down the breathing. Indeed, connectivity between olfactory and limbic structures is unique, the olfactory bulb directly projecting to limbic areas (Kay & Freeman, 1998; Soudry et al., 2011; Zhou et al., 2021). Consequently, odors are experienced primarily in terms of emotions (Bensafi et al., 2002; Herz, 2016). Among them, pleasant odors have a particular status. Indeed, pleasant odors slow down respiration rate and cardiac activity (Bensafi, 2002; Masaoka et al., 2012) while unpleasant ones induce rapid and superficial breathing (Masaoka et al., 2005). Similarly, only pleasant odors (*vs* unpleasant) are able to improve mood, and to decrease anxiety and pain unpleasantness (Villemure et al., 2003). However, the concept of pleasant smell should be treated with caution, as a concept specific to each individual, because evaluation of odor pleasantness varies drastically between people (Mantel et al., 2019).

While aromachology or olfactotherapy uses the potency of pleasant odors to influence mood, physiology, and behavior (Angelucci et al., 2014; Herz, 2009), little is known about how it works, nor whether the effects are specific to odorant stimuli. In this paper, our aim was to test the hypothesis that pleasant odors can specifically appease the body and the mind, with a multidisciplinary approach combining the analysis of subjective feelings of relaxation (questionnaires), physiological signals (heart rate and respiration), and brain activity (EEG). Through the privileged access of odors to the limbic system, its impact on respiration regime, and the influence of breathing on cerebral activity, a personally pleasant odor could *i)* enhance the subjective feeling of relaxation, *ii)* slow down body rhythms, and *iii)* facilitate entrainment of brain activity by respiration. Moreover, we assumed that the olfactory modality should be more effective than any other sensory one in inducing such effects. We tested this assumption by comparing the effects of a personally pleasant odor to a personally pleasant music on psychological, physiological and neuronal responses.

## 2 RESULTS

Because evaluation of a sensory stimulus pleasantness varies drastically between people, we made each participant choose their favorite smell and music during a first session (Fig.1A). As evidenced on Figure 2, the selected stimuli were all pleasant (pleasantness > 0.5) with a pleasantness of 0.91 (±0.07, [0.88, 0.93]; mean ± standard deviation, 95% confidence interval) for odors and of 0.93 (±0.07, [0.90, 0.95]) for music. Wilcoxon test did not reveal any statistical difference between pleasantness ratings of odors and music (W = 139, *p* = 0.15).

**Figure 1:**
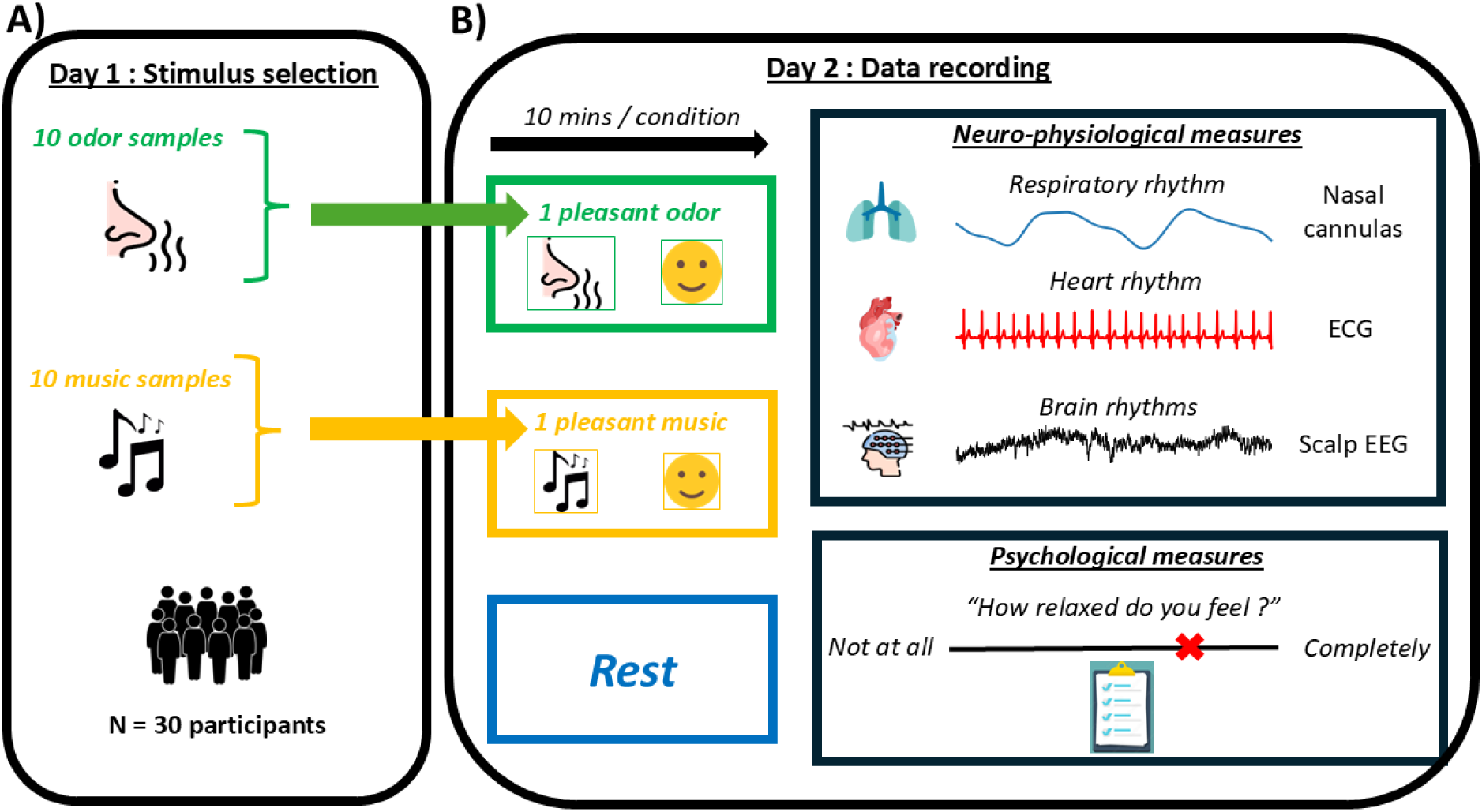
Experimental design. Thirty participants came for two sessions conducted on two different days: the first (A) aimed at selecting the most pleasant odor and the most pleasant music among a pre- selection of 10 that would be used in the second session (B), which aimed at recording physiological, neuronal and psychological data.

**Figure 2:**
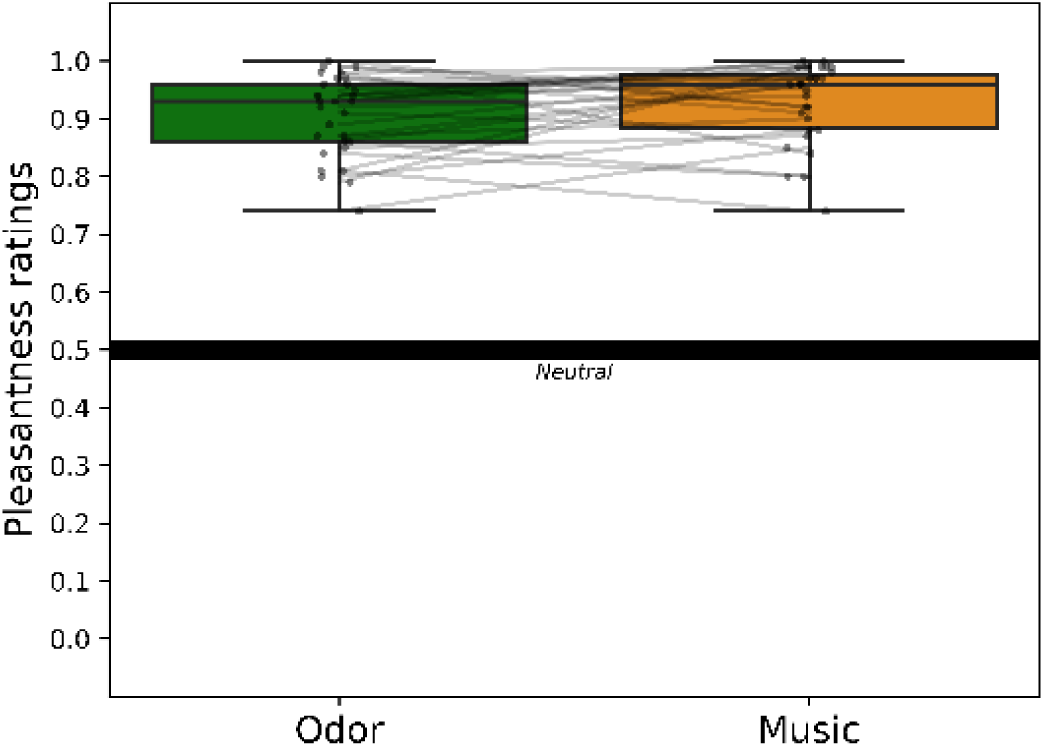
Odors and Music pleasantness ratings. The distribution of data is displayed as boxplots (minimum, first quartile, median, third quartile, maximum). Individual data points are superimposed on the plots, with lines connecting those from the same participant.

One day after session 1 (Fig.1B), a second session took place where the participant was equipped with sensors for body and brain rhythms recording: electrodes for cardiac activity, nasal cannulas for respiratory activity, EEG scalp electrodes for brain activity. The participant was then comfortably seated in front of a table on which were installed either an audio speaker or an articulated arm holding an odor-containing vial according to the block condition (music or odor respectively). The session consisted of three blocks of 10-minutes recordings during which participants were instructed to rest: a baseline block first, during which no stimulus was presented, followed by two blocks of sensory stimulation, odor or music, randomized and separated by a 15-minute break. During each block, physiological and EEG signals were recorded. After each block, participants evaluated, through paper questionnaires 1) the perceived intensity of relaxation at that time, and 2) the perceived arousal at that time, both on a continuous scale going from “not at all” to “completely”.

### 2.1 Neither odor nor music influenced subjective feeling of relaxation; both modified subjective feeling of arousal

Subjective feeling of relaxation was evaluated through the question “How relaxed do you feel?” that the participants had to quote on a continuous 10 centimeters paper scale going from “Not at all”, to “Completely” and values were normalized from 0 to 1. As evidenced in Figure 3A, subjective feeling of relaxation was of 0.64 (± 0.20, CI95: [0.56, 0.72]; mean ± standard deviation, CI95) after the baseline condition, 0.72 (± 0.17, [0.66, 0.78]) after the music condition, and 0.75 (± 0.12, [0.71, 0.80]) after the odor condition. Despite a tendency for both music and odor conditions to increase relaxation, no significant effect of experimental conditions on subjective relaxation was observed (W = 0.07, *p* = 0.12). Because the music selected by the participants had different tempi and because tempo plays an important role in inducing relaxation (Bernardi et al., 2006), we analyzed the correlation between tempo and subjective relaxation (Supplementary Figure 1A). Pearson correlation did not reveal a significant relationship between tempo and subjective relaxation (*r* = 0.26, *p* = 0.17; *r*² = 0.05, *p* = 0.24). Thus, neither odor nor music, whatever its tempo, significantly influenced subjective feeling of relaxation.

**Figure 3:**
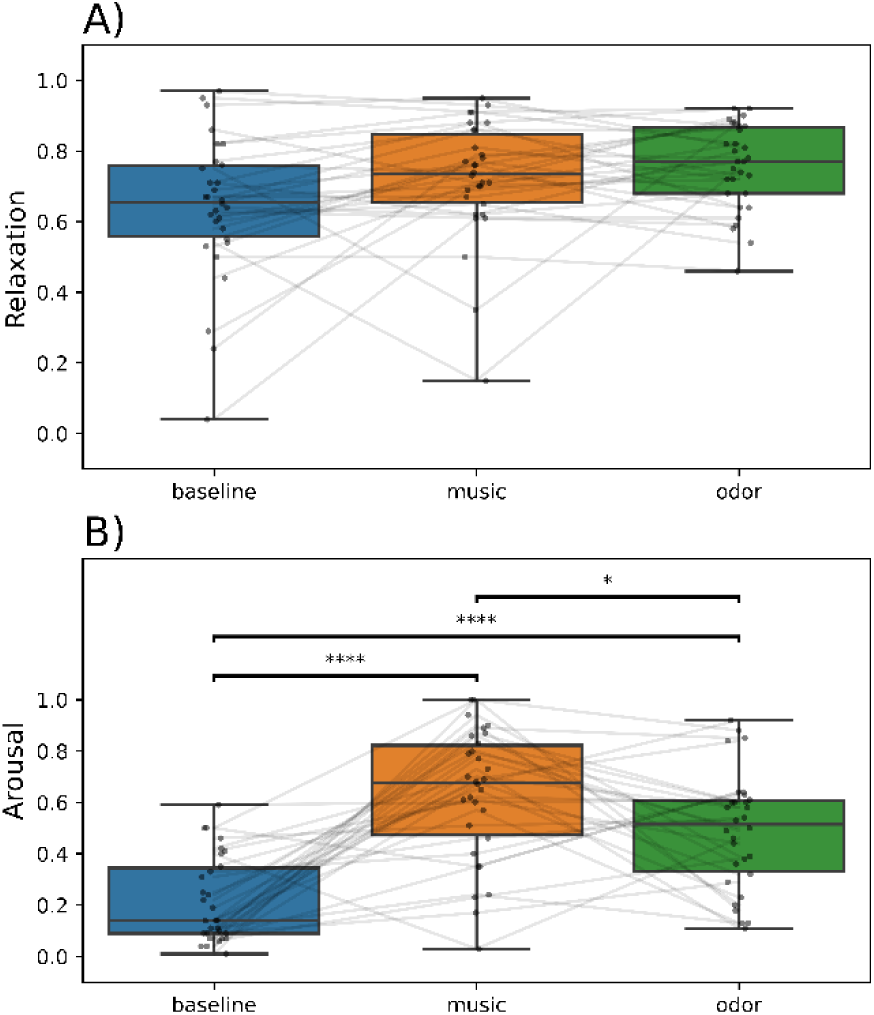
Effects of experimental conditions on measures of (A) relaxation and (B) arousal. The distribution of data is displayed as boxplots (minimum, first quartile, median, third quartile, maximum). Individual data points are superimposed on the plots, with lines connecting those from the same participant. *, p < 0.05; ***, p < 0.001.

Subjective feeling of arousal was explored through the question “How aroused do you feel?” using the same process and values were normalized from 0 to 1. As revealed in Figure 3B, subjective arousal was increased both during music and odor conditions. Subjective arousal significantly differed between experimental conditions (Friedman test, W = 0.58, *p* < 0.001) with both music (0.63 ± 0.26, [0.53, 0.73]) and odor (0.49 ± 0.23, [0.40, 0.57]) conditions being significantly different from baseline (0.22 ± 0.17, [0.16, 0.28], *p* < 0.0001). Therefore, both odor and music were perceived as arousing, even if this was more pronounced for music than for odor. Furthermore, no significant relationship between tempo of music and subjective arousal was found (*r* = -0.005, *p* = 0.98, *r*² = 0.001, *p* = 0.89; Supp. Fig.1B).

### 2.2 Pleasant smells, but not pleasant music, slowdown body rhythms

Body rhythms like respiratory and cardiac activities were recorded during the whole duration of the experiment.

#### 2.2.1 Respiratory activity

Respiratory signal was recorded through nasal cannulas connected to a pressure sensor capturing nasal airflow variations (see Methods). A homemade algorithm allowed detection of respiratory timestamps such as transitions from exhalation to inhalation and from inhalation to exhalation for each respiratory cycle. These detections allowed a cycle-by-cycle computing of respiratory features. For each experimental block and participant, we measured the median respiratory cycle duration (1/frequency), the median inspiratory volume, median expiratory volume and total median cycle volume.

Respiratory cycle durations were significantly different between conditions (Friedman test, W = 0.29, *p* < 0.001; Figure 4A). Respiratory cycles were shorter for music condition (4.14 ± 1.29 s, [3.66, 4.63]) and longer for odor condition (5.41 ± 2.06 s, [4.64, 6.18]) than baseline (4.71 ± 1.43 s, [4.18, 5.24]); *p’s* < 0.01). Same trends were observed for inhalation (Friedman test, W = 0.52, *p* < 0.001) and exhalation (Friedman test, W = 0.23, *p* < 0.001) durations that were both different between conditions (not shown). Indeed, inhalation durations were shorter for music condition (1.59 ± 0.52 s, [1.40, 1.79]) and longer for odor condition (2.20 ± 0.93 s, [1.85; 2.55]) than baseline (1.81 ± 0.54 s, [1.61, 2.01]). Similarly, exhalation durations were shorter for music condition (2.49 ± 0.83 s, [2.18, 2.80]) and longer for odor condition (3.16 ± 1.24 s, [2.70, 3.62]) than baseline (2.86 ± 0.98 s, [2.49, 3.23]). No significant relationship between tempo and respiratory cycle duration was revealed (Spearman correlation test, r = 0.019, p = 0.92; r² = 0.012, p = 0.57).

**Figure 4:**
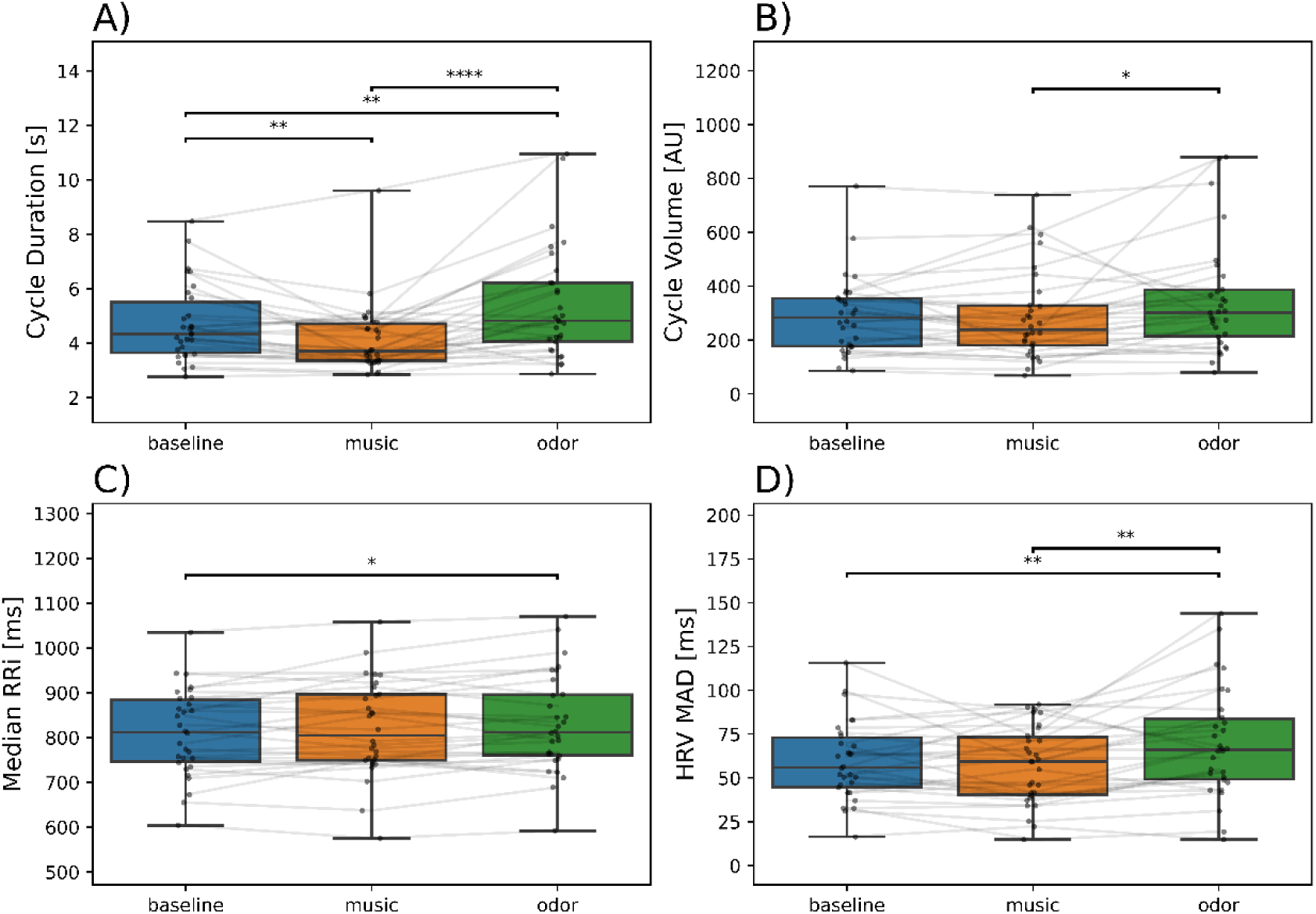
Effects of experimental condition on (A) respiratory cycle duration, (B) respiratory cycle volume, (C) median RRI from cardiac signal, and (D) HRV MAD from cardiac signal. The distribution of data is displayed as boxplots (minimum, first quartile, median, third quartile, maximum). Individual data points are superimposed on the plots, with lines connecting those from the same participant. *, p < 0.05; **, p < 0.01; ***, p < 0.001. AU, arbitrary unit.

Respiratory volumes significantly differed between conditions for inspiratory (Friedman test, W = 0.25, *p* < 0.001; not shown) and total respiratory cycle (Friedman test, W = 0.11, *p* = 0.04; Figure 4B). They were higher for odor condition compared to baseline (*p*’s < 0.05). No other *post-hoc* comparisons were significant (*p*’s > 0.05). Expiratory volume did not significantly differ between conditions (*p* = 0.08; not shown).

Overall, these results showed that a pleasant music speeded up breathing, whatever its tempo, while a pleasant odor slowed it down and increased its total volume.

#### 2.2.2 Cardiac activity

Cardiac signal was recorded using an electrocardiogram (ECG, see Methods) *via* three electrodes placed on the anterior surface of the right wrist, the left wrist, and in the lower abdominal region at the left iliac fossa. From the cardiac signal, *physio* (Ghibaudo et al., 2023), a python coded toolbox was used to detect R peaks and R-R intervals (RRI) were computed. RRI representing the time duration between consecutive heartbeats, median value of the RRIs was computed as a proxy of heart rate.

As shown in Figure 4C, median RRI significantly differed between experimental conditions (*F*(2,58) = 4.19, *p* = 0.02), with median RRI being significantly higher in odor condition (830.18 ± 108.55 ms, [789.65, 870.71]) than in baseline (811 ± 97.10 ms, [774.74, 847.26]; *p* < 0.05). No difference was observed with music condition compared to baseline (821.08 ± 106.35 ms, [781.37, 860.79]; *p* > 0.05). As for respiratory cycle duration, tempo could modulate heart rate according to its rhythm. However, no significant correlation was found between median RRI and tempi (*r* = 0.205, *p* = 0.28; *r*² = 0.03, *p* = 0.34; Supp. Fig.1B). Thus, these results showed that only pleasant odors increased median RRI, in other words decreased heart rate. Music did not induce variations of median RRI, whatever its tempo.

Median Absolute Deviation (MAD) of the computed RRI was used to extract Heart Rate Variability (HRV), which was used as a proxy of vagal tone. HRV MAD (Figure 4D) significantly differed between conditions (*F* (2,58) = 7.76, *p* = 0.001) with odor condition (69.56 ± 31.01 ms, [57.98, 81.14]) being significantly different from baseline (59.55 ± 22 ms, [51.19, 67.91]; *p* < 0.01). No difference was observed with music condition compared to baseline (57.28 ± 21.98 s, [49.07, 65.49]; *p* > 0.05). Again, no significant correlation was found between HRV MAD and tempi (*r* = 0.02, *p* = 0.92; *r*² = 0, *p* = 0.93; Supp. Fig.1C,). Thus, these results showed that only pleasant odor increased heart rate variability. Music did not induce variations of HRV MAD, whatever its tempo.

### 2.3 Pleasant smells, but not pleasant music, enhanced the coupling between respiratory and EEG activities

Our aim was to test whether a pleasant stimulus could influence the coupling between brain activity and respiration. We thus recorded scalp EEG activity, *via* 32 active EEG electrodes positioned according to the international 10-20 system using actiCAP nap (actiCAP Brain Products GmbH, Gilching, Germany), with the ground electrode placed in the mid-frontal region.

From the respiratory signal, we extracted the respiratory characteristics (Inspiration to Expiration transition points IE, Expiration to Inspiration transition EI points, inspiratory and expiratory amplitudes, and inspiratory, expiratory, and total volumes), on a cycle-by-cycle basis (see Methods). Using the *physio* toolbox (Ghibaudo et al., 2023), we rescaled the brain signal time basis based on the timestamps of respiratory phase transitions (EI and IE timestamps), allowing the EEG signal to be segmented as one neural epoch per respiratory cycle, that could be then averaged to obtain the mean EEG signal along respiratory cycle. The strength of the modulation of EEG activity by respiration (“Respiration-modulated EEG activity”) was extracted by measuring the amplitude of mean EEG signal (maximum value – minimum value) for each electrode, experimental condition, and participant. We then used cluster-based permutations tests to compare extracted values during music and odor conditions to those obtained in baseline (see Methods). Finally, only electrodes with significant different signals (stimulus condition *vs* baseline) were topographically represented as yellow spots (Figure 5).

**Figure 5:**
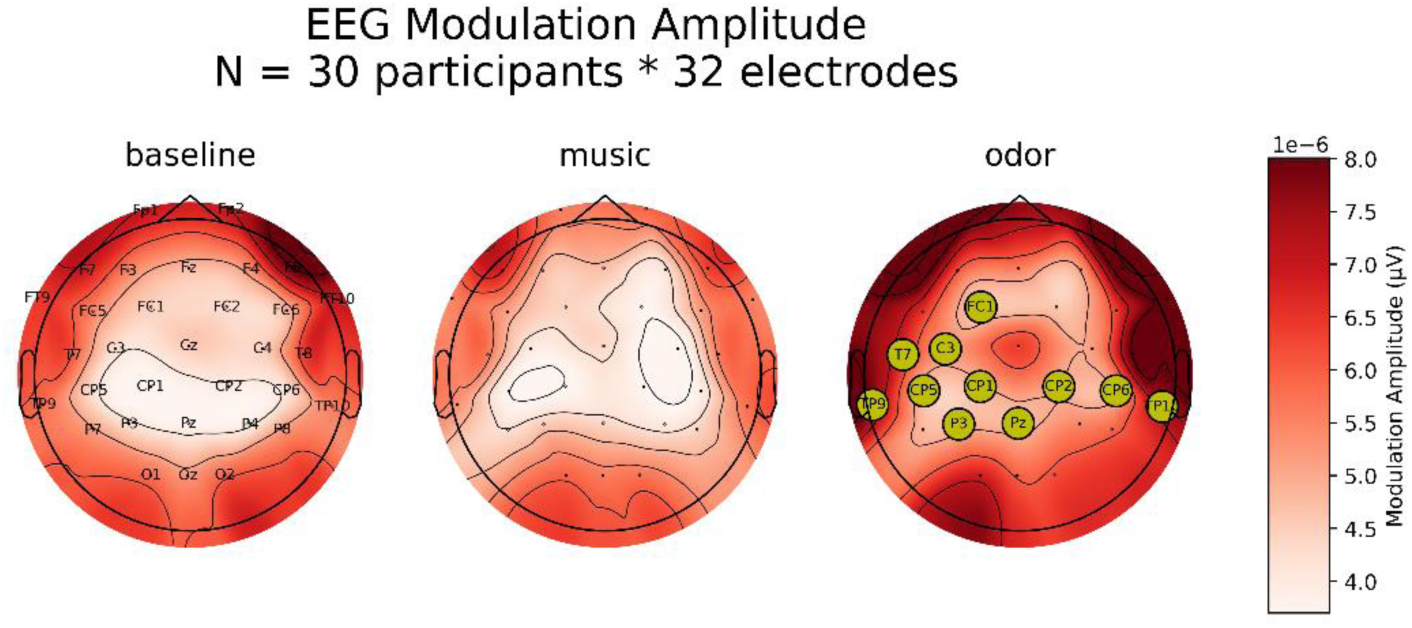
Respiration-modulated EEG activity induced by music and odors. Amplitude of modulation of the EEG signal by respiratory phase was obtained for each participant, electrode, and experimental condition. Averaged values are topographically represented for the three experimental blocks. The EEG modulation by the respiratory phase was significantly increased in comparison to baseline (p < 0.05) at electrodes displayed by yellow dots.

As evidenced by such topographic representation (Figure 5), amplitude of the respiration-related EEG activity significantly increased (*p* < 0.05) in a bilateral temporo-central cluster only during odor condition, music condition being not different from baseline.

We then tested whether such a respiratory drive could be modulated by the tempo of the different music. Cluster-based permutation tests comparing each condition of tempo to baseline did not allow evidencing an effect of music, whatever the tempo (Supplementary Fig.2).

Finally, since we observed that pleasant music and odors modified both respiratory cycle duration and volume, we investigated which of these parameters was more efficient in driving EEG activity. We explored the correlations between (A) duration and (B) volume of respiratory cycle and the amplitude of respiratory-modulated EEG activity (Figure 6). We observed that the amplitude of respiration-modulated EEG activity was significantly correlated with respiratory cycle duration (Figure 6A; *r* = 0.78, *p* < 0.0001) and with respiratory cycle volume (Figure 6B; *r* = 0.38, *p* < 0.001). The poorer correlation with volume compared to duration is explained by the fact that amplitude of respiratory- modulated EEG activity was not correlated with the respiratory cycle amplitude (Figure 6C; *r* = -0.007, *p* > 0.05). Probably involvement of emotions, relaxation, intéroception. But protocol was not design to test these hypothesis.

**Figure 6:**
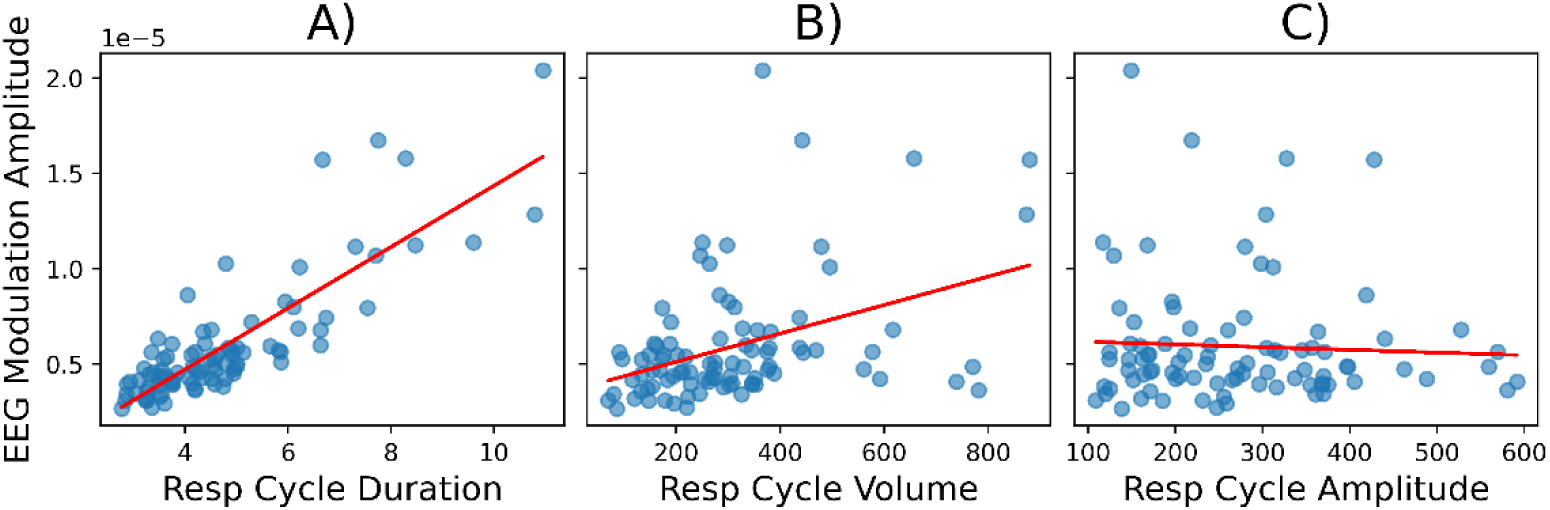
Correlation between respiratory features and respiration-modulated EEG activity. Average respiration-modulated EEG for each participant and experimental condition is represented according to A) the corresponding average respiratory cycle duration, B) the corresponding average respiratory cycle volume, C) the corresponding average respiratory cycle amplitude. A red line indicates the regression line. Resp, respiratory.

Overall, these results suggest that pleasant odor was more prone to induce a respiration-related activity in the brain than pleasant music, even if the music tempo was slow, and all the more so as it slows down the breathing rhythm.

## 3 DISCUSSION

The aim of the current study was to test the key role of olfaction in improving relaxation by comparing the effects of pleasant odors to those of pleasant music from three complementary angles: psychological, physiological and neuronal. Psychologically, we found for both odor and music a tendency to improve the subjective relaxation feeling, and a significant effect to increase arousal. Physiologically, we evidenced a soothing effect specific to odors: pleasant odor, but not pleasant music, slowed down respiration, increased inspiratory volume, decreased heart rate, and increased HRV. From the neuronal point of view, pleasant odor, but not pleasant music, increased the amplitude of the respiration-related EEG activity in a bilateral temporo-central region. Our study reveals that, rather than a general impact of stimulus pleasantness, the effects we observed are a peculiar contribution of odorants. We show for the first time that a personally pleasant odor is a sensory stimulus of choice to promote harmonization between body and brain rhythms.

### Pleasant odors favor the coupling between body and brain rhythms: the key role of olfactomotor response

Several research groups have already reported the powerful effect of odors on breathing and autonomic responses (Alaoui-Ismaïli et al., 1997; Royet et al., 2003). Pleasant odors, particularly, slow down breathing (Bensafi et al., 2003; Ferdenzi et al., 2015) and cardiac frequency (Bensafi, 2002a). Interestingly, an odor, that is not pleasant by itself, can nevertheless slow breathing frequency and increase amplitude if it has been previously associated with a pleasant taste (Yeomans & Prescott, 2016). Even when a participant is instructed to mentally imagine a smell, imagery of pleasant odors involves larger sniffs than imagery of unpleasant odors (Bensafi et al., 2003). In an episodic memory task, participants breathed in deeper and longer for pleasant odors than for unpleasant ones (Saive et al., 2014). Similarly, retrieval of a positive autobiographical memory triggered by personally pleasant odor is associated with a slower and deeper breathing compared to the perception of control odors evoking no memory (Masaoka et al., 2012). All these observations tend to show that an odorant stimulus triggers an olfactomotor response mainly characterized by breathing modification.

The breathing modification, here a slowdown, elicits first an autonomic response consisting in a heart rate variability increase. This could be due to the well-known cardio-respiratory coupling whose main consequence is that a respiratory rate decrease induces an increase in amplitude of respiratory sinus arrhythmia (RSA). RSA is a form of cardio-respiratory coupling characterized by variations in cardio- inhibitory vagal parasympathetic tone during the respiratory cycle (Berntson et al., 1993; Eckberg, 1983; Porges, 1986). This tone decreases during inhalation and increases during exhalation, leading to heart rate oscillations that are synchronized with respiration (Berntson et al., 1993; Menuet et al., 2020). Studies have demonstrated that the amplitude of these oscillations increases as respiratory rate decreases (Hayano et al., 1994; Hirsch & Bishop, 1981; Laude et al., 1993). Thus, it is very likely that the HRV increase we observed with odor-induced respiratory rate slowdown could be mostly explained by an increase in RSA amplitude. It is interesting to note that some theories link the withdrawal of RSA with anxiety (Campbell & Wisco, 2021), suggesting a possible association between increased RSA and a state of relaxation, although these ideas are highly debated.

We also evidenced that brain exhibits a greater respiration-related activity as respiration slows and deepens. The respiratory rhythm has been observed, both in rodents and humans, to influence brain oscillations across various brain regions through the impact of breathing-related airflows on olfactory receptor cells (Bagur et al., 2021; Hammer et al., 2021; Herrero et al., 2018; Ito et al., 2014; Juventin et al., 2022; Karalis & Sirota, 2022; Kluger et al., 2021; Tort et al., 2018; Yanovsky et al., 2014; Zelano et al., 2016). We previously showed in rodents that the slow and deep respiratory regime, characteristic of the rest state, is optimal for respiration to drive the brain (Girin et al., 2021; Juventin et al., 2022). Here, we complete this finding by showing in humans that a personally pleasant odor, through the olfactomotor response it provokes, can induce a slow rhythm in a large temporo-central network. Such slow brain rhythms offer sufficiently long temporal windows to recruit large neural assemblies, thus allowing the coordination of long-distance networks, as the ones involved in default mode activity of the brain (Buzsáki, 2010; Buzsáki & Watson, 2012; Raichle & Snyder, 2007). This raises the possibility that olfactomotor responses triggered by pleasant odors could facilitate the engagement of the brain in its default mode, thus promoting brain and body states of recovery (Beissner et al., 2013).

Our assumption is that most of the autonomic and cerebral effects we observed are a consequence of the olfactomotor response, as illustrated in Figure 7. Indeed, odors being experienced primarily in terms of emotions (Bensafi et al., 2002b; Herz, 2016; Royet et al., 2000), a pleasant odor first evokes an emotional response (“I like it”) which triggers an olfactomotor response (“I breathe it slower and deeper”), which in turn elicits both autonomic (heart rate decrease) and brain (respiration-modulated EEG activity) responses (Figure 7).

**Figure 7:**
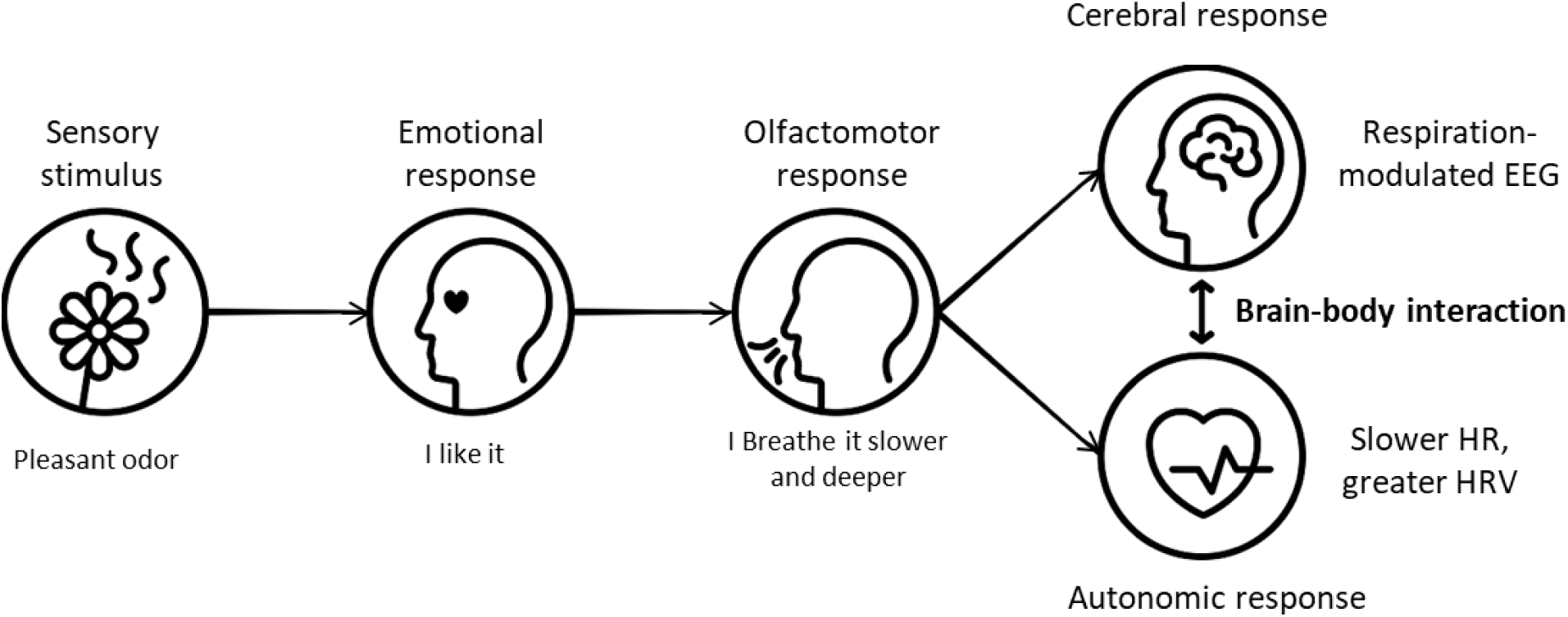
Hypothetical events sequence explaining the specific effects of pleasant odors on brain- body interaction.

That could explain the large temporo-central network of respiration-modulated EEG activity we observed under pleasant odor condition (Figure 5). This network could testify to the involvement of premotor, motor and somatosensory cortices, which are involved in volitional and attentive breathing (Herrero et al., 2018). Our protocol and EEG method do not allow us to exclude the involvement of regions related to emotions or interoception.

### Music pleasantness is not a relevant parameter for inducing coupling between body and brain rhythms

Conversely, we observed that a pleasant music, even with slow tempo, accelerated breathing rate and had no effect on cardiac activity. Music increased the subjective feeling of arousal more than pleasant odors did. This suggests that pleasant music tend to induce a physiological arousal while pleasant odors induce a physiological appeasement. This confirms what was described before that, compared with silence, music increases heart and respiratory frequencies, that are higher during pleasant than unpleasant music listening (Koelsch & Jäncke, 2015; Orini et al., 2010). However, other studies reported that certain types of music, especially those with slow tempi, slow down both cardiac and breathing rates (Bernardi et al., 2006). Recently, Baccarani et al. (Baccarani et al., 2023) reported that, during a stress recovery period, music elicited cardiac modifications. This discrepancy with our results could be due to different factors. For example, previous work found differential effects of tempi on the physiology on musician *vs* non-musician participants (Bernardi et al., 2006). Among plausible influencing factors, the listening context is probably one of the most contributing (Linnemann et al., 2015).

### Pleasant odors do not improve the subjective feeling of relaxation: a conflicting effect

We observed that, contrary to some previous findings, neither odors nor music significantly altered the subjective perception of relaxation despite a tendency to improve it (Davis & Thaut, 1989; Herz, 2016; Masaoka et al., 2012; Matsunaga et al., 2011; Wolfe et al., 2002). Given the extensive literature on the subject (Gard et al., 2014; Jerath et al., 2015; Sevoz-Couche & Laborde, 2022; Zaccaro et al., 2018), we expected that a slower respiratory rate, as that evoked by odors, would also be associated with a perception of increased relaxation. However, in our experiment, the subjective measure of relaxation did not correlate with what is generally considered as objective measures, *i.e.* breathing rate decrease and HRV increase (Zaccaro et al., 2018). Zaccaro et al. have previously noted “limited evidence” regarding the relationship between physiological and psychological outcomes in the context of slow breathing techniques and anxiety (Zaccaro et al., 2018). One possible reason for this lack of effect could be that participants were already in a state of calmness when arriving, as indicated by the relatively high baseline level of subjective relaxation (∼0.65). Another possibility is that our single question about the present feeling of relaxation could be not sufficient to assess such a complex feeling and a more in-depth questionnaire could provide a more accurate measurement and reduce the variability inherent in evaluating this state. Most importantly, behavioral and physiological outcomes of odors vary enormously among participants depending on many factors such as culture, experience, mood, expectation, and many others (for a review, see Herz, 2009). Several groups emphasized the importance of the notion of “personally pleasant” in the use of odorant stimuli (Herz, 2016; Masaoka et al., 2012; Matsunaga et al., 2011; Royet et al., 2000). We considered this parameter foremost here and this is one of the strengths of our study. Nevertheless, this could also be a limitation because, contrary to the pleasantness of the odor we fixed as high, the arousal power was left variable. In such conditions, a pleasant odor could vary from very poorly to very much arousing. Since arousal has major impact on autonomic nervous system (Azarbarzin et al., 2014; Bach et al., 2010; Bensafi, 2002; Wang et al., 2018; Wascher, 2021), it is not surprising that we observed no increase in the subjective feeling of relaxation, despite a real autonomic response increase.

## Conclusion

We demonstrated that the positive emotion evoked by a personally pleasant smell is sufficient to evoke an olfactomotor response, which, by slowing breathing, synchronizes respiration, fluctuations of heart rate and brain activity. Such a coupling between brain and body rhythms cannot be achieved by listening to a pleasant music, at least in our experimental conditions. Being easily achieved non-invasively, the use of odors (scents, perfumes, essential oils…) constitutes a potent avenue to improve bodily and cerebral state and thus mental state.

## Acknowledgments

This work was supported by the French National Research Agency (ANR-22-CE37-0014, BreathSmellRelax), and the Roudnitska Foundation. We express our special thanks to perfumer Jean-Charles Sommerard for the creation of the odor samples. We also thank Christelle Daude,Belkacem Messaoudi, Thibaut Woog and Hervé Hugueney for their technical assistance.

## Author contributions

Conceptualization and Methodology: MT, VG, JG, NB, JP; Supervision: NB; Experiments: MT, VG; Funding acquisition: NB; Software: VG, JG, SG; Writing – original draft: VG; Writing – review & editing: VG, MT, JG, NB, JP

## Supplemental information

**Supplementary Table 1:** Dilution table of odorant solutions. Odorants are provided by Jean-Charles Sommerard (Sevessence)

| Odorant | Dilution (% stock solution) |
| --- | --- |
| Blue ocean | 50 |
| Cotton flower | 20 |
| Olive leaf | 50 |
| Oxygen | 20 |
| Peach Lavender | 20 |
| Rose | 50 |
| Spiced orange blossom | 20 |
| Spiced wood | 10 |
| Spring floral | 50 |
| Vanilla | 75 |

**Supplementary Table 2:** Categories of music evaluated by participants.

| **Music genre** | **Reference** | **Average tempo (bpm)** |
| --- | --- | --- |
| Classical | Sir Neville Marriner - Was mir behagt, ist nur die muntre Jagd, BWV 208: IX. Schafe können sicher weiden (Orch. Marriner) (1713) | 55 |
| Electronic | Janji - Heroes Tonight (2015) | 128 |
| French variety | Isabelle Boulay – Parle-moi (2000) | 81 |
| Hard rock | AC/DC - Kick You When You Down (2020) | 110 |
| Jazz | Herbie Hancock - Speak Like a Child (1968) | 122 |
| Metal | Aephanemer - Path Of The Wolf (2019) | 134 |
| Pop | Miley Cyrus - Adore You (2013) | 60 |
| Rap #1 | Freeze Corleone - Freeze Raël (2020) | 68 |
| Rap #2 | Luv Resval - AZNVR (2023) | 108 |
| Raga | Zia Mohiuddin Dagar – Dhrupad, Raga Yaman (1991) | 56 |

bpm: beats per minute.

**Figure S1:**
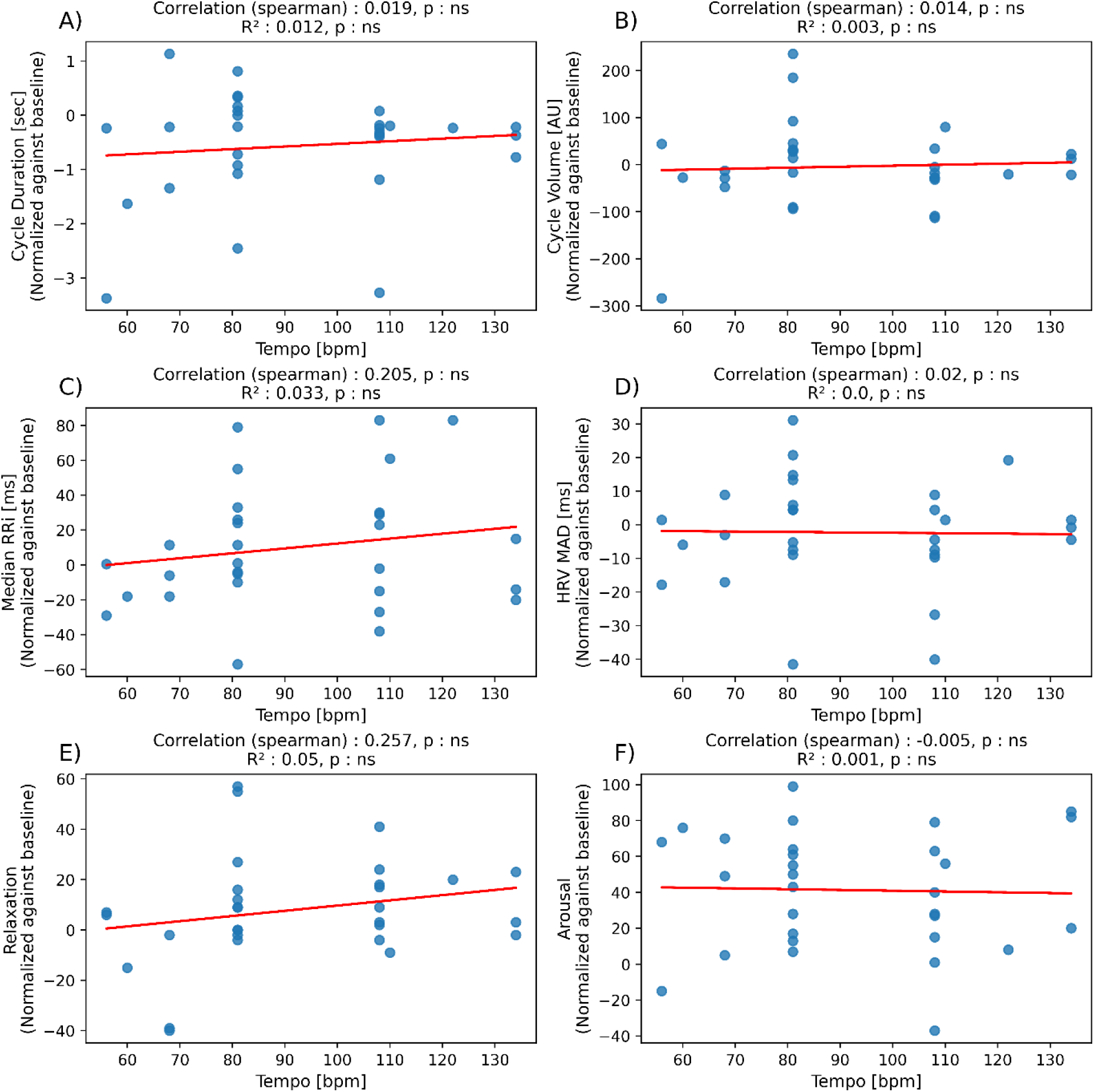
Effects of music tempi on physiological and subjective markers of relaxation (**A)** Respiratory cycle duration, (**B)** Respiratory total cycle volume, (**C)** Median value of RR interval, (**D)** Median Absolute Deviation of RR interval (HRV MAD), (**E)** Subjective relaxation according to music tempo, and (**F)** Subjective arousal. According to its chosen music, each participant would have listened to music with a particular tempo. These latter are presented in abscissa while values obtained for various metrics are presented in ordinate. Values of metrics computed from music block have been normalized by those obtained during baseline block (music data – baseline data). A red line indicates the regression line. Statistics are presented in title of each subplot: spearman correlation coefficient, regression coefficient, and interpretation of *p*-values (ns: non-significant, *: *p* < 0.05).

**Figure S2:**
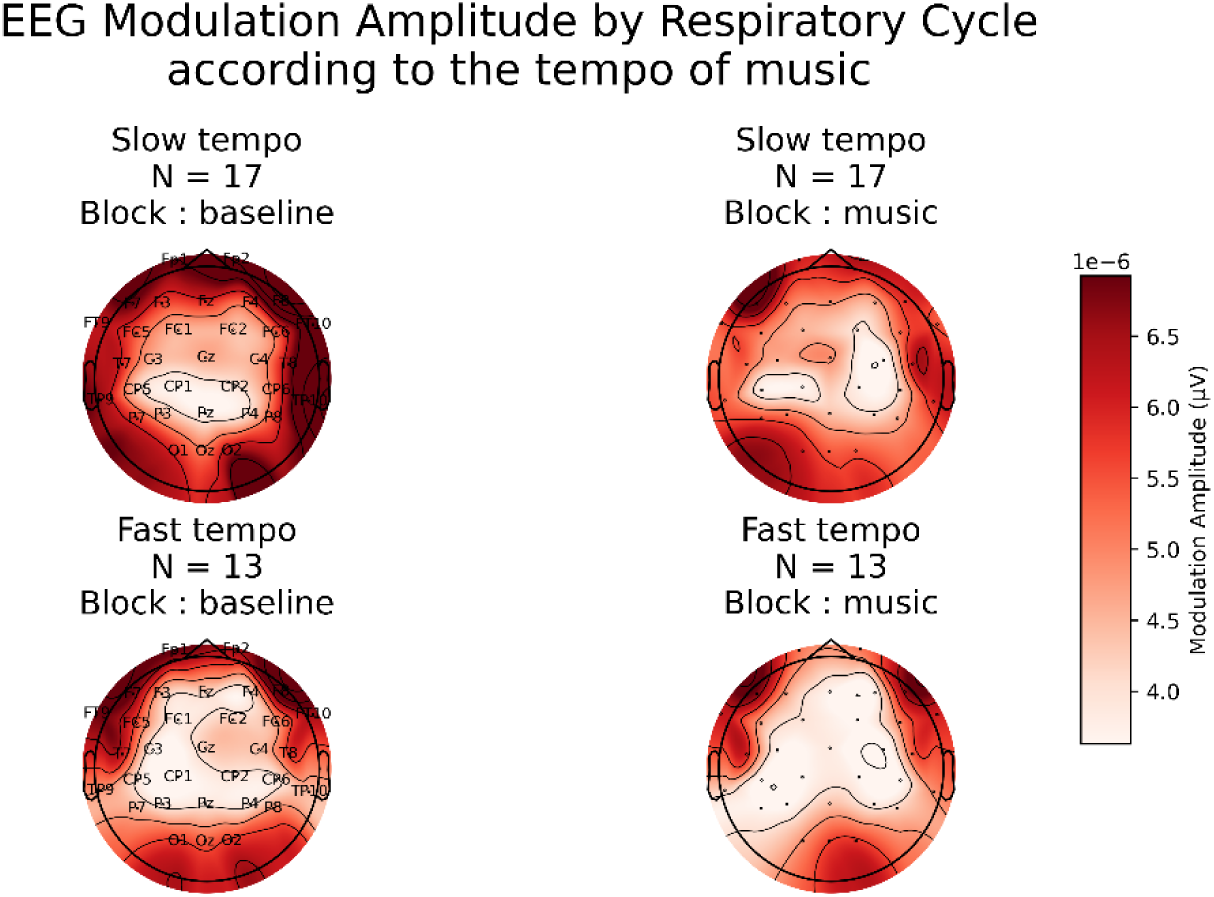
Effects of music tempi on EEG Modulation by respiratory cycle. Amplitude of modulation of the EEG signal by respiratory phase was obtained for each participant, electrode, for baseline (left) and music (right) condition. According to its chosen music, each participant would have listened to music with a particular tempo. Averaged values are topographically represented for the baseline and music blocks (left and right panels respectively) but split according to the tempo of the music chosen by the participants, Slow (<90 bpm; n = 17) and Fast (>90 bpm; n = 13) on upper and lower panels, respectively. The use of cluster-based permutations tests to compare music against baseline data of the two populations (tempi) did not allow evidencing an effect of tempo.

## 4 MATERIALS AND METHODS

### 4.1 Participants and Ethics Statement

Thirty participants (16 women, 14 men, aged 27.3 ± 9.5 years) took part in the study and attended two sessions at the Lyon Neuroscience Research Center (Registration number DPO service: 2-20125). Inclusion criteria were as follows: aged between 18 and 60 years, covered by social security, and willing to participate in the study. Exclusion criteria were as follows: pregnant, in labor, or breastfeeding women; individuals deprived of liberty by judicial or administrative decision; those with known cardiovascular or respiratory disorders; those with known olfactory, neurosensory, or psychiatric disorders. This study was approved on May 17, 2022, by an Ethics Committee, according to French regulations for biomedical experiments with healthy volunteers [CPP IDF3, ID RCB: 2021-A03077- 34]. Participants were informed in advance of the study in a clear and fair manner, provided written informed consent and were paid for their participation

### 4.2 Data and software

#### 4.2.1 Psychological measures

Psychological measure of relaxation was evaluated through the question “How relaxed do you feel?” that the participants had to quote on a continuous 10 centimeters paper scale going from “Not at all”, to “Completely”. Furthermore, psychological arousal was explored through the question “How aroused do you feel?” using the same process. Values were normalized from 0 to 1.

#### 4.2.2 Physiological measures

Respiratory signal was recorded through nasal cannulas connected to a pressure sensor (Sensortechnics GmbH, Puchheim, Germany) capturing nasal airflow variations. Cardiac signal was recorded using an electrocardiogram (ECG, Brain Products GmbH, Gilching, Germany) *via* three electrodes placed on the anterior surface of the right wrist, the left wrist, and in the lower abdominal region at the left iliac fossa. Both respiratory and ECG signals were acquired using a DC amplifier (actiCHamp Plus Brain Products GmbH, Gilching, Germany) at a sampling rate of 1000 Hz.

#### 4.2.3 Electroencephalographic activity

Electroencephalographic (EEG) activity was recorded through scalp EEG using the same amplifier and sampling frequency as for ECG and respiratory signals (see 2.2.2), *via* 32 active EEG electrodes positioned according to the international 10-20 system using actiCAP nap (actiCAP Brain Products GmbH, Gilching, Germany). The ground electrode was placed in the mid-frontal region. Impedances were kept below 50 kΩ by applying conductive gel to the scalp.

### 4.3 Stimuli and apparatus

#### 4.3.1 Odors

Ten complex and presumed pleasant odors were composed by a perfumer (Sevessence, Dardilly, France, SIRET: 75348599400036). They were titled Blue Ocean, Cotton Flower, Olive Leaf, Oxygen, Peach Lavender, Rose, Spiced Orange Blossom, Spiced Wood, Spring Floral, Vanilla. Fifteen-ml brown vials were filled three-quarters with polypropylene beads soaked in 1 mL of odorant solution diluted in isopropyl myristate (Hyteck Aroma-Zone, Paris, France). The dilution levels were adjusted for each perfume to ensure they were not overly intense while remaining easily perceptible for several tens of minutes. Supplementary Table 1 gives the dilution performed for each perfume. For delivery, the vials were held by a homemade articulated arm, 10 cm from the participants’ nose.

#### 4.3.2 Music pieces

Ten music pieces were preselected from the favorite genres of the French population (Supplementary Table 2). These genres included Classical music, Electronic, French variety, Hard rock, Jazz, Metal, Pop, Rap, and World music (raga was chosen). The selected music pieces were transformed into instrumental versions using Audacity software version 3.5.1 (https://www.audacityteam.org/), to prevent potential cognitive artifacts due to reciting lyrics in the mind. Average tempo of each sample was extracted through FL Studio software version 21 (https://www.image-line.com/) and confirmed by an experimented musician.

### 4.4 Procedure

The experimental protocol consisted of two sessions conducted on two different days: the first aimed at selecting the most pleasant odor and the most pleasant music that would be used in the second session, which aimed at recording physiological, neuronal and psychological data. The experimental protocol is schematically represented in **Erreur ! Source du renvoi introuvable.**.

#### 4.4.1 Session 1: Selection of individual most pleasant Odor and Music

In the first session, the participants rated the odors and music pieces in two different ways: an absolute rating followed by a relative rating. The absolute rating consisted in evaluating the pleasantness of each stimulus independently on a continuous scale going from extremely unpleasant to extremely pleasant, with a central bar indicating neutrality. Based on this, the three highest-rated stimuli were selected for the relative rating. These were then rated relatively to each other (*i.e.*, their ratings could be compared) according to their pleasantness. The odor and music rated with the highest pleasantness were selected for Session 2. Obtained values were normalized from 0 to 1 (0 for “extremely unpleasant” to 1 for “extremely pleasant”).

#### 4.4.2 Session 2: Recording Psychological, Physiological and Neuronal data in specific sensory environments

Second session took place 1 day after the first. The participant was asked to sit comfortably on a chair while keeping eyes open, and once equipped with sensors (EEG, ECG, nasal cannulas), was positioned in front of a desk, on which a mechanical arm was installed to keep an odorant vial close to the participant’s nose. A speaker (Essentielb SB70 Portable Speaker, Boulanger, Lille, France) was also placed on the desk one meter away. Three blocks of 10-minutes recordings were initiated during which participants were instructed to rest: a baseline block first, during which no stimulus was presented, followed by two blocks of sensory stimulation (odor or music), randomized and separated by a 15- minute break. After each block, participants were asked to evaluate, through paper questionnaires 1) the perceived intensity of relaxation at the moment, and 2) the perceived arousal at the moment, both on a continuous scale going from “not at all” to “completely”. Values were normalized from 0 to 1.

### 4.5 Data analysis

#### 4.5.1 Physiological signals

All analysis was performed using a homemade Python toolbox (Ghibaudo et al., 2023). We describe briefly here the main steps of processing.

*Respiratory cycles* were extracted as follows (Figure 8). First, the respiratory signal was detrended and smoothed using a low-pass filter with a 7 Hz Bessel type. Then, Inspiration-Expiration Transitions (IE) and Expiration-Inspiration Transitions (EI) were detected when respiratory signal raised and crossed its median level, and decayed and crossed its median level, respectively. Based on these detected timestamps, respiratory cycle characteristics, such as durations, amplitudes, and volumes, were computed cycle by cycle. Finally, aberrant respiratory cycles, resulting from faulty detection or artifacts, were statistically removed to ensure data accuracy and reliability according to Ghibaudo et al., 2023.

**Figure 8:**
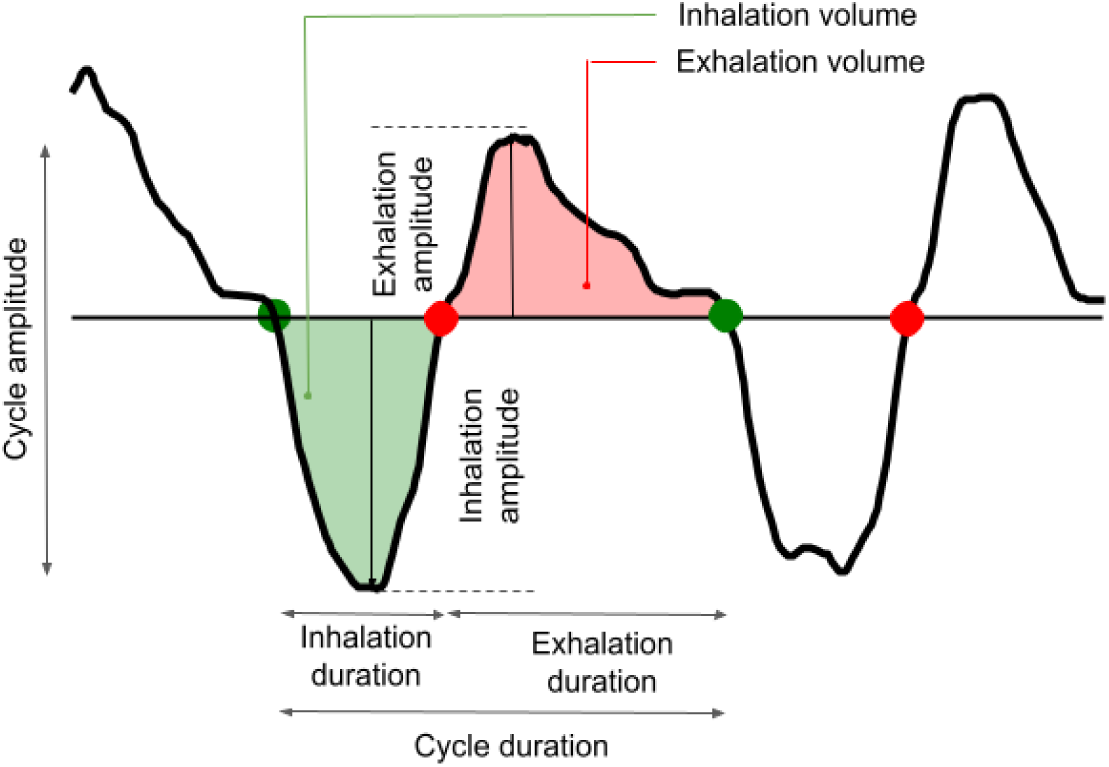
Detection of respiratory timestamps and calculation of respiratory features. The *physio* toolbox allows processing of respiratory traces (black) and thus detection of respiratory timestamps such as transitions from exhalation to inhalation (green dots) and from inhalation to exhalation (red dots) for each respiratory cycle. These detections allow the computing of respiratory features for each respiratory cycle, based on duration and amplitude of each phase (inspiration, expiration, cycle), that allows inferring the variations of respiratory regime. Modified from Ghibaudo et al., 2023.

*ECG signal* of each participant and experimental condition was first preprocessed by centered and band- pass filtered using a Bessel filter (bandpass 5 to 45 Hz). This step aimed to improve the signal-to-noise ratio to isolate the R peaks of the signal by removing other characteristic ECG waves considered as noise. Next, R peaks were detected with a minimum separation of 400 milliseconds. Then, R-R intervals (RRI) were computed, representing the time duration between consecutive heartbeats. Median value of the RRIs was computed as a proxy of heart rate. Median Absolute Deviation (MAD) of these computed RRI was used to extract Heart Rate Variability.

#### 4.5.2 EEG signals

##### 4.5.2.1 Preprocessing

Signal recorded by each electrode was re-referenced to the common average. Slow drifts were removed through linear detrending, and resulting traces were centered by subtracting the median value itself. Signal was band-stopped (50 Hz, Notch filter for line noise removal) and band-passed (0.05 to 200 Hz, Butterworth type, order 6). Independent Component Analysis was applied to exclude ocular artifact components (blinks and ocular-motor movements, characterized by high-amplitude phasic components, especially in the fronto-polar region). This process was performed using the MNE library, version 1.4.2 (Gramfort et al., 2013). Artifacted channels were manually removed. Finally, movement artifacts and the concerned respiratory epochs were removed based on the criteria of large statistical deviations (> 4.5 median absolute deviations from median baseline level) of gamma power.

##### 4.5.2.2 Measurement of respiration-related EEG activity

The respiratory characteristics (IE points, EI points, inspiratory and expiratory amplitudes, and inspiratory, expiratory, and total volumes), extracted on a cycle-by-cycle basis, was utilized to achieve a cyclical deformation of the preprocessed brain signal based on the respiratory phase. This step was automated by the *deform_traces_to_cycle_template()* function coded in the physio toolbox (Ghibaudo et al., 2023) that rescales the brain signal time basis based on the timestamps of respiratory phase transitions (EI and IE timestamps) through linear interpolation. This rescaled signal was segmented to provide one neural epoch per respiratory cycle with a common time vector (the respiratory phase) that was averaged to obtain the mean EEG signal along respiratory cycle. Then the strength of the modulation of EEG activity by respiration (“Respiration-modulated EEG activity”) was extracted by measuring the amplitude of mean EEG signal (maximum value – minimum value) for each electrode, experimental condition, and participant. This process is schematically represented in Figure 9.

**Figure 9:**
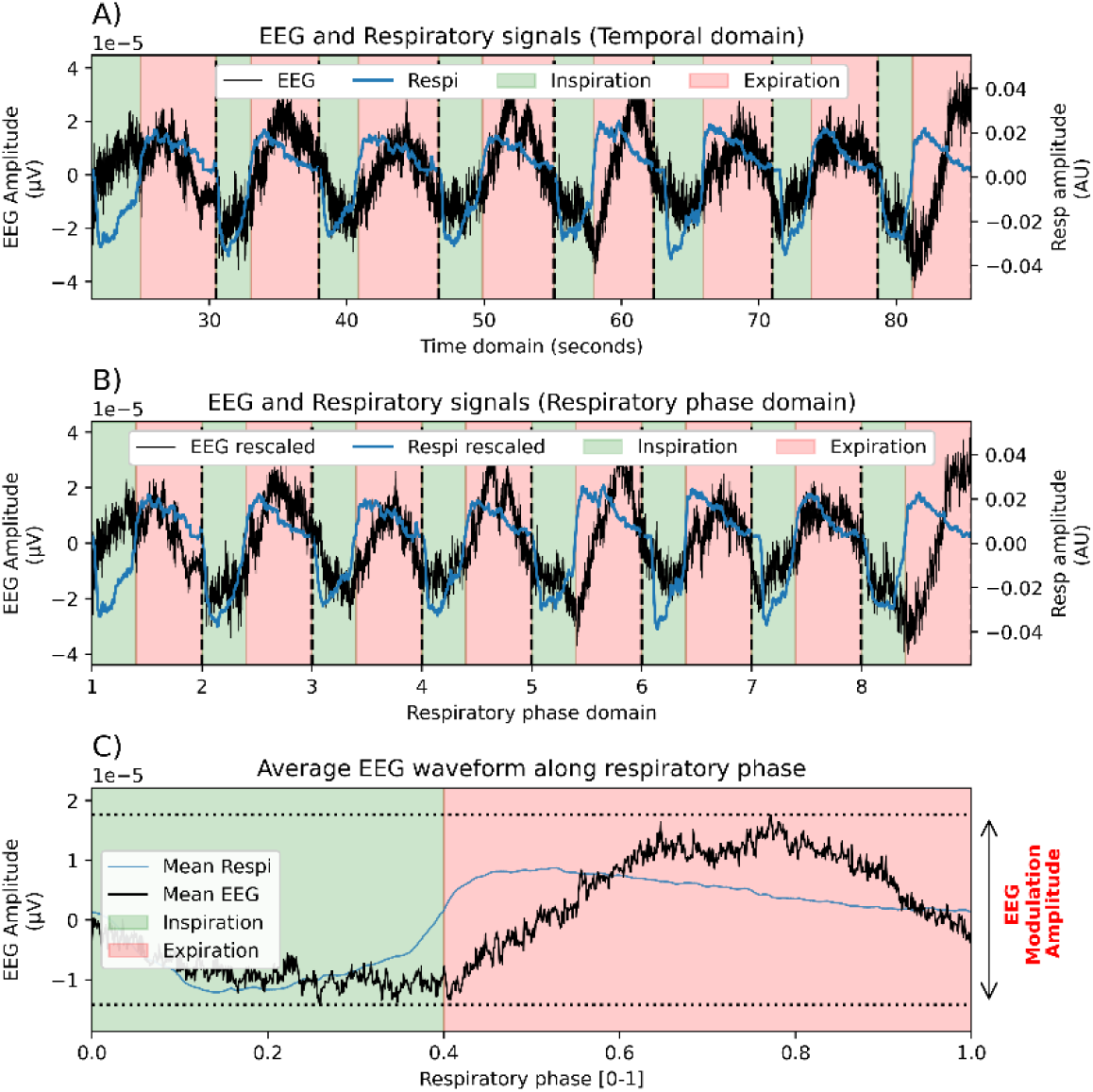
Pipeline of measurement of respiration-related EEG activity. **A)** An example of EEG time series is presented in black and the corresponding respiratory time series (nasal airflow) is presented in blue. Inspiratory and expiratory epochs deduced from respiratory feature detection are figured in green and red spans, respectively. Black vertical dashed lines indicate the separation between two successive respiratory cycles. Note the variability of duration of the eight successive respiratory epochs/cycles, that would prevent averaging of respiratory-based EEG epochs. **B)** Time series are rescaled into respiratory phase-based series through linear interpolation of traces between detected respiratory timestamps (IE and EI transitions). Therefore, each respiration-based EEG epoch has the same length and can be averaged across epochs/cycles. **C)** Averaged EEG cyclical dynamic along respiratory phase is displayed in black. The total amplitude of this trace can be computed (max – min) to extract a strength of modulation of EEG amplitude by respiration: “EEG Modulation Amplitude”. AU, arbitrary unit; Respi, respiration.

### 4.6 Statistics

Except for cluster-based permutation statistics used for analyzing EEG results, we used statistical tests recruited from *pingouin* (Vallat, 2018), a Python coded toolbox dedicated to statistics, in order to compare the three experimental conditions: baseline, odor, music. Data were considered significantly different if *p*-value result was < 0.05.

#### 4.6.1 Respiratory Cycle Duration, Cycle Volume, Median RRi, HRV MAD, Relaxation, Arousal

Psychological and physiological metrics were obtained with a sample size of N = 30, repeatedly (within-design). We firstly assessed for the sphericity and normality of the data, deduced from Mauchly and Shapiro-Wilk test, respectively. When data presented a default of sphericity or normality, we used a non-parametric within-design test allowing for comparison of three conditions: Friedman test. This was the case for all metrics except for Median RRI and HRV MAD for which we used parametric within-design test: repeated-measure analyses of variance (ANOVA). If the effect of experimental conditions was significantly different, we extended explorations by using two-sided pairwise *post-hoc* tests: *t*-tests or Wilcoxon test according to the sphericity and normality of data. *p*-values were adjusted for multiple comparison with Holm–Bonferroni method.

#### 4.6.2 Respiration-related EEG activity

The EEG Modulation Amplitude was obtained for each participant, electrode, and experimental condition. Conserving topographical relationship of the obtained values allowed for the computing of cluster-based permutations tests to compare music and odor extracted values to those obtained in baseline. To do this, we used the *mne.stats.permutation_cluster_1samp_test()* function coded in the toolbox MNE (Gramfort et al., 2013). This function allows for non-parametric cluster-level paired *t*-test as described in Maris and Oostenveld (Maris & Oostenveld, 2007). We provided to the function a 30 (participants) * 32 (electrodes) sized matrix with the difference between odor and baseline conditions. The same process was used to compare music to baseline condition. The threshold parameter of the function was set to none (default), meaning that the cluster-forming threshold was chosen automatically based on a *p*-value of 0.05 for the given number of observations. The number of permutations was set to the default: 1024. The tail parameter was set to 0, meaning that the statistic is computed on both sides of the null distributions.

#### 4.6.3 Correlations and regressions

Statistical interactions between metrics were explored through computation of quantitative statistics with Pearson correlations and linear regressions by using *scipy.stats.pearsonr* and *scipy.stats.linregress* functions from *scipy* toolbox (Virtanen et al., 2020).

## Code availability statement

All analysis scripts used for this study are available at this link: https://github.com/ValentinGhibaudo/Scripts_Odor_Music_Brain_Body

## Abbreviations

BL: Baseline
BMRQ: Barcelona Music Reward Questionnaire
CPP: Comité de Protection des Personnes (Ethics Committee)
ECG: Electrocardiogram
EEG: Electroencephalogram
HRV: Heart Rate Variability
ICA: Independent Component Analysis
MAD: Median Absolute Deviation
OAS: Odor Awareness Scale
RSA: Respiratory Sinus Arrhythmia
ANS: Autonomic Nervous System

## Bibliography

Alaoui-Ismaïli, O., Robin, O., Rada, H., Dittmar, A., & Vernet-Maury, E. (1997). Basic emotions evoked by odorants : Comparison between autonomic responses and self-evaluation. Physiology & Behavior, 62(4), 713-720. 10.1016/s0031-9384(97)90016-0

Angelucci, F. L., Silva, V. V., Dal Pizzol, C., Spir, L. G., Praes, C. E. O., & Maibach, H. (2014). Physiological effect of olfactory stimuli inhalation in humans : An overview. International Journal of Cosmetic Science, 36(2), 117-123. 10.1111/ics.12096

Azarbarzin, A., Ostrowski, M., Hanly, P., & Younes, M. (2014). Relationship between Arousal Intensity and Heart Rate Response to Arousal. Sleep, 37(4), 645-653. 10.5665/sleep.3560

Baccarani, A., Donnadieu, S., Pellissier, S., & Brochard, R. (2023). Relaxing effects of music and odors on physiological recovery after cognitive stress and unexpected absence of multisensory benefit. Psychophysiology, 60(7), e14251. 10.1111/psyp.14251

Bach, D. R., Friston, K. J., & Dolan, R. J. (2010). Analytic measures for quantification of arousal from spontaneous skin conductance fluctuations. International Journal of Psychophysiology, 76(1), 52-55. 10.1016/j.ijpsycho.2010.01.011

Bagur, S., Lefort, J. M., Lacroix, M. M., de Lavilléon, G., Herry, C., Chouvaeff, M., Billand, C., Geoffroy, H., & Benchenane, K. (2021). Breathing-driven prefrontal oscillations regulate maintenance of conditioned-fear evoked freezing independently of initiation. Nature Communications, 12(1), 2605. 10.1038/s41467-021-22798-6

Beissner, F., Meissner, K., Bär, K.-J., & Napadow, V. (2013). The autonomic brain : An activation likelihood estimation meta-analysis for central processing of autonomic function. The Journal of Neuroscience: The Official Journal of the Society for Neuroscience, 33(25), 10503-10511. 10.1523/JNEUROSCI.1103-13.2013

Bensafi, M. (2002). Autonomic Nervous System Responses to Odours : The Role of Pleasantness and Arousal. Chemical Senses, 27(8), 703-709. 10.1093/chemse/27.8.703

Bensafi, M., Porter, J., Pouliot, S., Mainland, J., Johnson, B., Zelano, C., Young, N., Bremner, E., Aframian, D., Khan, R., & Sobel, N. (2003). Olfactomotor activity during imagery mimics that during perception. Nature Neuroscience, 6(11), 1142-1144. 10.1038/nn1145

Bensafi, M., Rouby, C., Farget, V., Bertrand, B., Vigouroux, M., & Holley, A. (2002a). Influence of affective and cognitive judgments on autonomic parameters during inhalation of pleasant and unpleasant odors in humans. Neuroscience Letters, 319(3), 162-166. 10.1016/s0304-3940(01)02572-1

Bensafi, M., Rouby, C., Farget, V., Bertrand, B., Vigouroux, M., & Holley, A. (2002b). Psychophysiological correlates of affects in human olfaction. Neurophysiologie Clinique/Clinical Neurophysiology, 32(5), 326-332. 10.1016/S0987-7053(02)00339-8

Bernardi, L., Porta, C., & Sleight, P. (2006). Cardiovascular, cerebrovascular, and respiratory changes induced by different types of music in musicians and non-musicians : The importance of silence. Heart, 92(4), 445-452. 10.1136/hrt.2005.064600

Berntson, G. G., Cacioppo, J. T., & Quigley, K. S. (1993). Respiratory sinus arrhythmia : Autonomic origins, physiological mechanisms, and psychophysiological implications. Psychophysiology, 30(2), 183-196. 10.1111/j.1469-8986.1993.tb01731.x

Boiten, F. A., Frijda, N. H., & Wientjes, C. J. (1994). Emotions and respiratory patterns : Review and critical analysis. International Journal of Psychophysiology: Official Journal of the International Organization of Psychophysiology, 17(2), 103-128. 10.1016/0167-8760(94)90027-2

Brown, R. P., & Gerbarg, P. L. (2009). Yoga breathing, meditation, and longevity. Annals of the New York Academy of Sciences, 1172, 54-62. 10.1111/j.1749-6632.2009.04394.x

Buzsáki, G. (2010). Neural syntax : Cell assemblies, synapsembles, and readers. Neuron, 68(3), Article 3.

Buzsáki, G., & Watson, B. O. (2012). Brain rhythms and neural syntax : Implications for efficient coding of cognitive content and neuropsychiatric disease. Dialogues in Clinical Neuroscience, 14(4), 345-367.

Campbell, A. A., & Wisco, B. E. (2021). Respiratory sinus arrhythmia reactivity in anxiety and posttraumatic stress disorder : A review of literature. Clinical Psychology Review, 87, 102034. 10.1016/j.cpr.2021.102034

Davis, W. B., & Thaut, M. H. (1989). The Influence of Preferred Relaxing Music on Measures of State Anxiety, Relaxation, and Physiological Responses 1. *Journal of Music Therapy*, *26*(4), 168-187. 10.1093/jmt/26.4.168

Dlouhy, B. J., Gehlbach, B. K., Kreple, C. J., Kawasaki, H., Oya, H., Buzza, C., Granner, M. A., Welsh, M. J., Howard, M. A., Wemmie, J. A., & Richerson, G. B. (2015). Breathing Inhibited When Seizures Spread to the Amygdala and upon Amygdala Stimulation. Journal of Neuroscience, 35(28), 10281-10289. 10.1523/JNEUROSCI.0888-15.2015

Eckberg, D. L. (1983). Human sinus arrhythmia as an index of vagal cardiac outflow. *Journal of Applied Physiology: Respiratory*, Environmental and Exercise Physiology, 54(4), 961-966. 10.1152/jappl.1983.54.4.961

Ferdenzi, C., Fournel, A., Thévenet, M., Coppin, G., & Bensafi, M. (2015). Viewing Olfactory Affective Responses Through the Sniff Prism : Effect of Perceptual Dimensions and Age on Olfactomotor Responses to Odors. Frontiers in Psychology, 6, 1776. 10.3389/fpsyg.2015.01776

Folschweiller, S., & Sauer, J.-F. (2021). Respiration-Driven Brain Oscillations in Emotional Cognition. Frontiers in Neural Circuits, 15. https://www.frontiersin.org/articles/10.3389/fncir.2021.761812

Fulwiler, C. E., & Saper, C. B. (1984). Subnuclear organization of the efferent connections of the parabrachial nucleus in the rat. Brain Research Reviews, 7(3), 229-259. 10.1016/0165-0173(84)90012-2

Gard, T., Noggle, J. J., Park, C. L., Vago, D. R., & Wilson, A. (2014). Potential self-regulatory mechanisms of yoga for psychological health. Frontiers in Human Neuroscience, 8, 770. 10.3389/fnhum.2014.00770

Ghibaudo, V., Granget, J., Dereli, M., Buonviso, N., & Garcia, S. (2023). A Unifying Method to Study Respiratory Sinus Arrhythmia Dynamics Implemented in a New Toolbox. eNeuro, 10(10). 10.1523/ENEURO.0197-23.2023

Girin, B., Juventin, M., Garcia, S., Lefèvre, L., Amat, C., Fourcaud-Trocmé, N., & Buonviso, N. (2021). The deep and slow breathing characterizing rest favors brain respiratory-drive. Scientific Reports, 11(1), Article 1. 10.1038/s41598-021-86525-3

Gramfort, A., Luessi, M., Larson, E., Engemann, D., Strohmeier, D., Brodbeck, C., Goj, R., Jas, M., Brooks, T., Parkkonen, L., & Hämäläinen, M. (2013). MEG and EEG data analysis with MNE-Python. Frontiers in Neuroscience, 7. https://www.frontiersin.org/articles/10.3389/fnins.2013.00267

Grosmaitre, X., Santarelli, L. C., Tan, J., Luo, M., & Ma, M. (2007). Dual functions of mammalian olfactory sensory neurons as odor detectors and mechanical sensors. Nature Neuroscience, 10(3), 348-354. 10.1038/nn1856

Hammer, M., Schwale, C., Brankačk, J., Draguhn, A., & Tort, A. B. L. (2021). Theta-gamma coupling during REM sleep depends on breathing rate. Sleep, 44(12), zsab189. 10.1093/sleep/zsab189

Harper, R. M., Frysinger, R. C., Trelease, R. B., & Marks, J. D. (1984). State-dependent alteration of respiratory cycle timing by stimulation of the central nucleus of the amygdala. Brain research, *306*(1-2), Article 1-2.

Hayano, J., S, M., M, S., A, O., K, T., & T, F. (1994). Effects of respiratory interval on vagal modulation of heart rate. The American Journal of Physiology, *267*(1 Pt 2). 10.1152/ajpheart.1994.267.1.H33

Hegoburu, C., Shionoya, K., Garcia, S., Messaoudi, B., Thévenet, M., & Mouly, A.-M. (2011). The RUB Cage : Respiration-Ultrasonic Vocalizations-Behavior Acquisition Setup for Assessing Emotional Memory in Rats. Frontiers in Behavioral Neuroscience, 5, 25. 10.3389/fnbeh.2011.00025

Herrero, J. L., Khuvis, S., Yeagle, E., Cerf, M., & Mehta, A. D. (2018). Breathing above the brain stem : Volitional control and attentional modulation in humans. Journal of Neurophysiology, 119(1), 145-159. 10.1152/jn.00551.2017

Herz, R. S. (2009). Aromatherapy Facts and Fictions : A Scientific Analysis of Olfactory Effects on Mood, Physiology and Behavior. International Journal of Neuroscience, 119(2), 263-290. 10.1080/00207450802333953

Herz, R. S. (2016). The Role of Odor-Evoked Memory in Psychological and Physiological Health. Brain Sciences, 6(3), 22. 10.3390/brainsci6030022

Hirsch, J. A., & Bishop, B. (1981). Respiratory sinus arrhythmia in humans : How breathing pattern modulates heart rate. The American Journal of Physiology, 241(4), H620–629. 10.1152/ajpheart.1981.241.4.H620

Ito, J., Roy, S., Liu, Y., Cao, Y., Fletcher, M., Lu, L., Boughter, J. D., Grün, S., & Heck, D. H. (2014). Whisker barrel cortex delta oscillations and gamma power in the awake mouse are linked to respiration. Nature Communications, 5. Scopus. 10.1038/ncomms4572

Jerath, R., Crawford, M. W., Barnes, V. A., & Harden, K. (2015). Self-regulation of breathing as a primary treatment for anxiety. Applied Psychophysiology and Biofeedback, 40(2), 107-115. 10.1007/s10484-015-9279-8

Juventin, M., Ghibaudo, V., Granget, J., Amat, C., Courtiol, E., & Buonviso, N. (2022). Respiratory influence on brain dynamics : The preponderant role of the nasal pathway and deep slow regime. Pflugers Archiv: European Journal of Physiology. 10.1007/s00424-022-02722-7

Juventin, M., Zbili, M., Fourcaud-Trocmé, N., Garcia, S., Buonviso, N., & Amat, C. (2023). Respiratory rhythm modulates membrane potential and spiking of non-olfactory neurons. Journal of Neurophysiology. 10.1152/jn.00487.2022

Karalis, N., & Sirota, A. (2022). Breathing coordinates cortico-hippocampal dynamics in mice during offline states. Nature Communications, 13(1), Article 1.

Kay, L. M., & Freeman, W. J. (1998). Bidirectional processing in the olfactory-limbic axis during olfactory behavior. Behavioral Neuroscience, 112(3), 541-553. 10.1037/0735-7044.112.3.541

Kluger, D. S., Balestrieri, E., Busch, N. A., & Gross, J. (2021). Respiration aligns perception with neural excitability. eLife, 10, e70907. 10.7554/eLife.70907

Koelsch, S., & Jäncke, L. (2015). Music and the heart. European Heart Journal, 36(44), 3043-3049. 10.1093/eurheartj/ehv430

Lacuey, N., Hampson, J. P., Harper, R. M., Miller, J. P., & Lhatoo, S. (2019). Limbic and paralimbic structures driving ictal central apnea. Neurology, 92(7), e655-e669. 10.1212/WNL.0000000000006920

Laude, D., Goldman, M., Escourrou, P., & Elghozi, J. L. (1993). Effect of breathing pattern on blood pressure and heart rate oscillations in humans. Clinical and Experimental Pharmacology & Physiology, 20(10), 619-626. 10.1111/j.1440-1681.1993.tb01643.x

Linnemann, A., Ditzen, B., Strahler, J., Doerr, J. M., & Nater, U. M. (2015). Music listening as a means of stress reduction in daily life. Psychoneuroendocrinology, 60, 82-90. 10.1016/j.psyneuen.2015.06.008

Mantel, M., Ferdenzi, C., Roy, J.-M., & Bensafi, M. (2019). Individual Differences as a Key Factor to Uncover the Neural Underpinnings of Hedonic and Social Functions of Human Olfaction : Current Findings from PET and fMRI Studies and Future Considerations. Brain Topography, 32(6), 977-986. 10.1007/s10548-019-00733-9

Maris, E., & Oostenveld, R. (2007). Nonparametric statistical testing of EEG- and MEG-data. Journal of Neuroscience Methods, 164(1), 177-190. 10.1016/j.jneumeth.2007.03.024

Masaoka, Y., & Homma, I. (2001). The effect of anticipatory anxiety on breathing and metabolism in humans. Respiration Physiology, 128(2), 171-177. 10.1016/S0034-5687(01)00278-X

Masaoka, Y., Koiwa, N., & Homma, I. (2005). Inspiratory phase-locked alpha oscillation in human olfaction : Source generators estimated by a dipole tracing method. The Journal of Physiology, 566(3), 979-997. 10.1113/jphysiol.2005.086124

Masaoka, Y., Sugiyama, H., Katayama, A., Kashiwagi, M., & Homma, I. (2012). Slow Breathing and Emotions Associated with Odor-Induced Autobiographical Memories. Chemical Senses, 37(4), 379-388. 10.1093/chemse/bjr120

Matsunaga, M., Isowa, T., Yamakawa, K., Kawanishi, Y., Tsuboi, H., Kaneko, H., Sadato, N., Oshida, A., Katayama, A., Kashiwagi, M., & Ohira, H. (2011). Psychological and physiological responses to odor-evoked autobiographic memory. Neuro Endocrinology Letters, 32(6), 774-780.

Menuet, C., Connelly, A. A., Bassi, J. K., Melo, M. R., Le, S., Kamar, J., Kumar, N. N., McDougall, S. J., McMullan, S., & Allen, A. M. (2020). PreBötzinger complex neurons drive respiratory modulation of blood pressure and heart rate. eLife, 9, e57288. 10.7554/eLife.57288

Orini, M., Bailón, R., Enk, R., Koelsch, S., Mainardi, L., & Laguna, P. (2010). A method for continuously assessing the autonomic response to music-induced emotions through HRV analysis. Medical & Biological Engineering & Computing, 48(5), 423-433. 10.1007/s11517-010-0592-3

Porges, S. W. (1986). Respiratory Sinus Arrhythmia : Physiological Basis, Quantitative Methods, and Clinical Implications. In P. Grossman, K. H. L. Janssen, & D. Vaitl (Éds.), Cardiorespiratory and Cardiosomatic Psychophysiology (p. 101-115). Springer US. 10.1007/978-1-4757-0360-3_7

Raichle, M. E., & Snyder, A. Z. (2007). A default mode of brain function : A brief history of an evolving idea. NeuroImage, 37(4), 1083-1090; discussion 1097-1099. 10.1016/j.neuroimage.2007.02.041

Rhone, A. E., Kovach, C. K., Harmata, G. I., Sullivan, A. W., Tranel, D., Ciliberto, M. A., Howard, M. A., Richerson, G. B., Steinschneider, M., Wemmie, J. A., & Dlouhy, B. J. (2020). A human amygdala site that inhibits respiration and elicits apnea in pediatric epilepsy. JCI Insight, 5(6), e134852, 134852. 10.1172/jci.insight.134852

Royet, J. P., Zald, D., Versace, R., Costes, N., Lavenne, F., Koenig, O., & Gervais, R. (2000). Emotional responses to pleasant and unpleasant olfactory, visual, and auditory stimuli : A positron emission tomography study. The Journal of Neuroscience: The Official Journal of the Society for Neuroscience, 20(20), 7752-7759. 10.1523/JNEUROSCI.20-20-07752.2000

Royet, J.-P., Plailly, J., Delon-Martin, C., Kareken, D. A., & Segebarth, C. (2003). fMRI of emotional responses to odors : Influence of hedonic valence and judgment, handedness, and gender. NeuroImage, 20(2), 713-728. 10.1016/S1053-8119(03)00388-4

Saive, A.-L., Royet, J.-P., Ravel, N., Thévenet, M., Garcia, S., & Plailly, J. (2014). A unique memory process modulated by emotion underpins successful odor recognition and episodic retrieval in humans. Frontiers in Behavioral Neuroscience, 8, 203. 10.3389/fnbeh.2014.00203

Sevoz-Couche, C., & Laborde, S. (2022). Heart rate variability and slow-paced breathing:when coherence meets resonance. Neuroscience & Biobehavioral Reviews, 135, 104576. 10.1016/j.neubiorev.2022.104576

Soudry, Y., Lemogne, C., Malinvaud, D., Consoli, S.-M., & Bonfils, P. (2011). Olfactory system and emotion : Common substrates. *European Annals of Otorhinolaryngology*, Head and Neck Diseases, 128(1), 18-23. 10.1016/j.anorl.2010.09.007

Tort, A. B. L., Brankačk, J., & Draguhn, A. (2018). Respiration-Entrained Brain Rhythms Are Global but Often Overlooked. Trends in Neurosciences, 41(4), 186-197. 10.1016/j.tins.2018.01.007

Vallat, R. (2018). Pingouin : Statistics in Python. Journal of Open Source Software, 3(31), 1026. 10.21105/joss.01026

Villemure, C., Slotnick, B. M., & Bushnell, M. C. (2003). Effects of odors on pain perception : Deciphering the roles of emotion and attention. Pain, *106*(1-2), 101-108. 10.1016/s0304-3959(03)00297-5

Virtanen, P., Gommers, R., Oliphant, T. E., Haberland, M., Reddy, T., Cournapeau, D., Burovski, E., Peterson, P., Weckesser, W., Bright, J., van der Walt, S. J., Brett, M., Wilson, J., Millman, K. J., Mayorov, N., Nelson, A. R. J., Jones, E., Kern, R., Larson, E., … van Mulbregt, P. (2020). SciPy 1.0 : Fundamental algorithms for scientific computing in Python. Nature Methods, 17(3), Article 3. 10.1038/s41592-019-0686-2

Wang, C.-A., Baird, T., Huang, J., Coutinho, J. D., Brien, D. C., & Munoz, D. P. (2018). Arousal Effects on Pupil Size, Heart Rate, and Skin Conductance in an Emotional Face Task. Frontiers in Neurology, 9. 10.3389/fneur.2018.01029

Wascher, C. A. F. (2021). Heart rate as a measure of emotional arousal in evolutionary biology. Philosophical Transactions of the Royal Society B: Biological Sciences, 376(1831), 20200479. 10.1098/rstb.2020.0479

Wolfe, D. E., O’Connell, A. S., & Waldon, E. G. (2002). Music for relaxation : A comparison of musicians and nonmusicians on ratings of selected musical recordings. Journal of Music Therapy, 39(1), 40-55. 10.1093/jmt/39.1.40

Yanovsky, Y., Ciatipis, M., Draguhn, A., Tort, A. B. L., & Brankačk, J. (2014). Slow Oscillations in the Mouse Hippocampus Entrained by Nasal Respiration. Journal of Neuroscience, 34(17), 5949-5964. 10.1523/JNEUROSCI.5287-13.2014

Yeomans, M. R., & Prescott, J. (2016). Smelling the goodness : Sniffing as a behavioral measure of learned odor hedonics. Journal of Experimental Psychology. Animal Learning and Cognition, 42(4), 391-400. 10.1037/xan0000120

Zaccaro, A., Piarulli, A., Laurino, M., Garbella, E., Menicucci, D., Neri, B., & Gemignani, A. (2018). How Breath-Control Can Change Your Life : A Systematic Review on Psycho-Physiological Correlates of Slow Breathing. Frontiers in Human Neuroscience, 12. https://www.frontiersin.org/article/10.3389/fnhum.2018.00353

Zelano, C., Jiang, H., Zhou, G., Arora, N., Schuele, S., Rosenow, J., & Gottfried, J. A. (2016). Nasal Respiration Entrains Human Limbic Oscillations and Modulates Cognitive Function. The Journal of Neuroscience, 36(49), 12448-12467. 10.1523/JNEUROSCI.2586-16.2016

Zhou, G., Olofsson, J. K., Koubeissi, M. Z., Menelaou, G., Rosenow, J., Schuele, S. U., Xu, P., Voss, J. L., Lane, G., & Zelano, C. (2021). Human hippocampal connectivity is stronger in olfaction than other sensory systems. Progress in Neurobiology, 201, 102027. 10.1016/j.pneurobio.2021.102027

